# Host-symbiont stress response to lack-of-sulfide in the giant ciliate mutualism

**DOI:** 10.1101/2021.07.27.454000

**Authors:** Salvador Espada-Hinojosa, Judith Drexel, Julia Kesting, Edwin Kniha, Iason Pifeas, Lukas Schuster, Jean-Marie Volland, Helena C. Zambalos, Monika Bright

## Abstract

The thiotrophic mutualism between the sulfur-oxidizing, chemoautotrophic (thiotrophic) bacterial ectosymbiont *Candidatus* Thiobius zoothamnicola and the giant ciliate *Zoothamnium niveum* thrives in a variety of shallow-water marine environments with highly fluctuating sulfide emission. To persist over time both partners must reproduce and ensure symbiont transmission prior cessation of sulfide, fueling the symbiont’s carbon fixation and host nourishment. We experimentally investigated the response of this mutualism to waning of sulfide. We found that colonies followed the r-strategy and released initially present but also newly produced macrozooids until death. A fraction of middle-sized longer-lived colonies were particularly proficient in producing and releasing swarmers. The symbionts on the colonies proliferated less and became larger and more rod-shaped under oxic conditions compared to symbionts from freshly collected colonies exposed to sulfide and oxygen. The symbiont monolayer was highly disturbed with epigrowth of other microbes and loss of symbionts that were subsequently found in the experimental seawater. We conclude that both partners’ response to cessation of sulfide emission was remarkably fast. The colony experienced death within two days but host reproduction through swarmers carrying the symbiont ensured the continuation of the association.

## Introduction

While aerobic eukaryotes die when exposed for extended times to one of the most dangerous poisons, hydrogen sulfide ([1]; ∑H_2_S i.e. sum of all forms of dissolved sulfide [2], hereafter called sulfide) most mutualistic associations between protist or animal hosts and sulfur-oxidizing chemoautotrophic (thiotrophic) bacteria depend on the presence of sulfide (see [3]). Thiotrophic symbionts share the need for reduced sulfur species (e.g. exclusively sulfide or additionally thiosulfate) and oxygen or alternative electron acceptors to gain energy for carbon fixation [3, 4]. The hosts provide sulfide and oxygen through uptake and transport to the symbionts or through specific behavior such as swimming in and out of vent fluid in shrimps, contraction/expansion behavior in colonial ciliates, or digging with the foot in some bivalves (see [3, 5, 6]). The hosts are, as far as is known, also capable of detoxifying sulfide (see [1]). In return, the symbionts nourish their hosts, which either lack a digestive tract and are entirely dependent on their symbionts, or supplement the nutrition provided by the symbionts by uptake of nutrients from the environment (see [3]).

Many habitats of these symbioses, however, are relatively short-lived, such as fast-spreading deep-sea hydrothermal vents, whale and wood falls depending on substrate size, and decaying seagrass debris (see [3]). In contrast to geothermally generated hydrogen sulfide as in vents (see [7]), biological sulfide production by sulfate-reducing bacteria ceases when organic material is depleted (see [8]). Upon changes in abiotic conditions, mobile animal hosts may migrate to more suitable habitats, e.g. stilbonematine nematodes, gutless oligochaetes, snails and bathymodiolin mussels (see [3]). Sessile hosts like siboglinid tubeworms [9] or peritrich ciliates [10–13], however, do not have this option.

To persist over generations, hosts reproduce primarily by releasing motile larvae into the pelagial. Regardless of their mobility as adults, larvae of bathymodiolin mussels, lucinid clams, and tubeworms disperse without their symbionts [14]. In these systems host reproduction and symbiont transmission are decoupled and the uptake of symbionts from a free-living population happens upon larval settlement. Experiments with some of these symbioses in oxic, non-sulfidic seawater showed that bathymodiolins and lucinids either lost their symbionts or highly reduced their density. Nevertheless, hosts could survive between one to five months until the end of experiments [15–18]. In contrast, experiments with tubeworms showed that their symbionts escaped upon host death [19]. The larvae of other hosts like vesicomyid and solemyid clams carry their symbionts (see [14]). Whether such hosts with vertically transmitted symbionts respond to sulfide starvation with loss of symbionts has not been studied. Moreover, it is not known whether sessile hosts of thiotrophic symbionts keep on reproducing under stress.

The symbiotic mutualism of the giant colonial ciliate *Zoothamnium niveum* (short *Zoothamnium*) and its thiotrophic gammaproteobacterial ectosymbiont, originally described as *Cand*. Thiobios zoothamnicoli but due to nomenclature regulations corrected to *Cand*. Thiobius zoothamnicola, ([20], short Thiobius) is a suitable model to study host-symbiont response to environmental stress and disturbance when sulfide ceases. In contrast to slow-growing and reproducing animal hosts, this is a fast-growing and fast reproducing, sessile ciliate [21] thriving on ephemeral sulfide-emitting surfaces in shallow-water environments such as wood, mangrove peat, decaying seagrass, and whale bones from tropical to temperate waters [22].

*Zoothamnium* colonies are composed of a stalk and alternate branches on which feeding microzooids, dividing terminal zooids, and macrozooids grow [11]. The dual partnership involves a single bacterial phylotype covering the host surface in a monolayer except for the lowest, senescent parts of the colonies. There, the symbionts get overgrown or replaced by other microbes [12,23,24]. The symbiont is rod-shaped except for the oral part of the microzooids on which they are more coccoid-shaped [12]. This phenotypic difference has been explained by the movement of cilia around the oral ciliature of feeding microzooids providing the symbionts with more balanced inorganic compounds for growth compared to all other cilia-free host parts [21].

The host colonies are sessile and can only ‘escape’ sulfide starvation through producing macrozooids and releasing them as swarmers [11, 12]. Experiments in custom-designed flow-through chambers creating a steady flow of oxic seawater mixed with sulfide showed that growth was highest at sulfide concentrations between 3 and 33 μmol L^-1^ [21]. Yet, *Zoothamnium* was able to grow also aposymbiotically into smaller colonies under oxic conditions without sulfide [21,25]. We do not know, however, how long large symbiotically grown colonies can survive under oxic conditions when sulfide ceases and what happens to their symbionts. Further it is unknown, whether the host releases swarmers carrying the symbionts as a response to oxic conditions and how long these swarmers survive in order to find a suitable habitat for settlement.

Here, we experimentally mimicked the waning of sulfide, a perturbation that is hypothesized to lead to stress and disturbance in this thiotrophic mutualism. We specifically asked how long freshly collected colonies and their swarmers survive under experimental oxic, non-sulfidic (short oxic) conditions. We also investigated if the release of macrozooids developed prior collection from submerged wood and the production of new macrozooids and their release as swarmers under oxic conditions is related to colony size. Further we studied the symbiont morphology, their frequency of dividing cells (FDC), as well as their density and coverage on the host surface, and the colonization of other microbes under oxic conditions using scanning electron microscopy (SEM) and fluorescence *in situ* hybridization (FISH). Given the lack of sulfide fueling symbiont carbon fixation, we hypothesized that the symbiont division should cease and consequently the monolayer on the host should be disrupted. To investigate whether the symbionts detach from the colony when exposed to prolonged lack-of-sulfide stress we implemented 16S rRNA Sanger sequencing to search for the symbiont in the incubation water.

## Material and Methods

### Ethic statement

No specific permissions were required for the listed locations as they are publicly accessible. Furthermore, we confirm that our field studies did not involve endangered or protected species.

### Sampling

*Zoothamnium niveum* colonies were collected from shallow, subtidal submerged woods at two locations in the Northern Adriatic Sea close to Piran, Slovenia: the estuarine canal Sv. Jernej (45°29’48.6”N, 13°35’57.0”E) and the mudflat in Strunjan (45°31’44.0”N, 13°36’13.2”E). Simultaneously, water samples were taken adjacent to the wood pieces and *in situ* temperature, salinity, and pH were measured using a Multi 340i sensor WTW (S1 Table). Wood pieces were transported in buckets filled with on-site seawater to the laboratory and maintained in flow-through aquaria until colonies were used for the experiments, from immediately after collection up to several days later. During maintenance about 250 mL of 1 mmol L^-1^ sulfide solution was added to each 50 L liter aquarium daily during which fresh seawater flow was stopped for a few hours. Each colony was cut off the wood with a MicroPoint™ Scissor and cleaned from debris by rinsing it in 0.2 μm filtered seawater prior the experimental procedure.

### Host response to oxic conditions in multi-well plates

60 colonies from each of the two collection sites Sv. Jernej and Strunjan, were placed individually into the wells of a multiwell plate filled each with 1 mL oxic, 0.2 μm filtered seawater. The number of macrozooids present on each colony was counted prior starting the experiment. Every 12 h, viability of colonies was assessed by their contraction/expansion behavior. Colonies that did not contract when being touched with a dissecting needle were considered dead. All swarmers released from each colony within 12 h time intervals were transferred into individual wells. Every 12 h about two-thirds of the water from each well was pooled for measurements of temperature, salinity, pH, and oxygen concentration (S2 Table). Oxygen was measured with a PreSenS Flow-through Cell FTC-PSt3. Wells were then filled up to 1 mL with fresh 0.2 μm filtered seawater. To estimate colony size, the number of branches was counted either after host death or at the end of the experiment (n = 85, only two colonies survived 96 h). Swarmers were mounted on glass slides and their body size was estimated using Leica DM2000 light microscope equipped with a Leica DFC295 camera and the image analysis software Gimp (GNU Image Manipulation Program) for Mac 2.8.

For statistical comparisons, 60 colonies from each location were divided into four batches (A-D), with 15 colonies each. The size of the swarmers was measured according to the timeframe of 12-h-interval observations they were released from the colony and the timeframe they were kept in the water swimming (batch A 0 h, B 24 h, C 48 h, D 72 h; S1 Fig). All time points were considered as the upper bound of the intervals. Statistical analyses were conducted in PAST 3.04 [26] and R [27]. Because Shapiro-Wilk tests showed deviations from normality for all parameters (counted branches taken as estimate of colony size, initial macrozooids per colony, total released swarmers per colony, and swarmer size) the Wilcoxon-Mann-Whitney test for equal medians was used for comparisons of the two locations. Mortalities of colonies and swarmers were estimated as the proportion of dead colonies/swarmers to the total number of colonies/swarmers used in the experiment. LT_50_ and their standard error estimates for colonies and swarmers were obtained by curve fitting of a binomial Generalized Linear Model (bGLM) with mortality rate as response and time as predictor, and the use of the R package MASS version 7.3-51.4. Goodness of fit was characterized with the Deviance (D^2^) = (Null Deviance-Residual Deviance)/(Null Deviance). Ordinary Least Squares (OLS) regression-models were used to depict the correlation between relevant magnitudes, e.g. colony size and released swarmers. The number of macrozooids developed during the experiment (Δ_M_) was calculated by subtracting the initial number of macrozooids present on colonies prior to the start of the experiment from sum of the released swarmers plus the macrozooids remaining on the colony at the end of the experiment. Δ_S_, the number of macrozooids produced and released as swarmers during the experiment, was calculated by the subtraction of the initial number of macrozooids from the number of released swarmers. A positive Δ_S_ value indicates additionally produced swarmers under oxic conditions during the experimental time frame, whereas a negative Δ_S_ value indicates remaining macrozooids on the respective colony that were not released anymore until the end of the experiment (S2 Fig).

### Symbiont response to oxic conditions in embryo dishes

In 2012, 2013, and 2014, sets of 15 to 20 freshly collected colonies from Sv. Jernej were put into embryo dishes, each kept completely filled with 0.2 μm filtered, oxic seawater and covered with glass plates to avoid evaporation for up to 72 h. At the time points 12, 24, 48, and 72 h viability of colonies was assessed as described above, and at most 3 live colonies were removed and fixed for SEM or divided into live and dead colonies and fixed for FISH (S1 Table). At each time point water was replaced with 0.2 μm filtered seawater (S2 Table). In 2013, additionally 15 to 21 mL of the incubation water pooled from the embryo dishes was taken after each time point and filtered through a 0.2 μm GTTP filter. One half of the filter was frozen in liquid nitrogen, the other half was fixed in 100% ethanol.

### Fluorescence *in situ* hybridization (FISH)

Colonies from the 2014 embryo dish experiments were fixed and stored in 100% ethanol at 4°C for 3 months. Colonies were embedded in LR-White resin and polymerization was performed in absence of oxygen at 41°C for three days. Semi-thin sections (1 μm) were cut on a Reichert Ultracut S microtome, placed in a drop of 20% acetone on chromium(III)potassium sulfate coated glass slides and were left to dry at 40°C. A total of 16 sections were placed on one slide with four spots of four sections each. To have a representative area of the colony on each slide, two slides per sample were used.

Hybridization was carried out as described in [24]. On each slide the symbiont-specific oligonucleotide probe ZNS196_mod [28] labeled with Cy3 together with a mix of EUB I, II, III (targeting most bacteria, [29,30]) and Arc 915 (targeting most archaea, [31]), all labeled with Cy5 to distinguish the symbiont from any other microbe, were used on two spots. Negative controls with nonsense probes (NON-EUB) labeled in both colors [32] were run on each slide on two different spots. In brief, applied probes were hybridized at 46°C for 3 hours in the dark, then rinsed in the washing buffer at 48°C for 15 min, stained with DAPI, washed with Milli Q, and mounted with Citifluor antifading solution. Sections were observed on a Zeiss Axio Imager M2 epifluorescence microscope and images were taken at 100x magnification with an AxioCam MRm, Zeiss using AxioVision Rel. 4.8. software. Composite pictures of entire colony sections were done with ICE software (Image Composite Editor 2.0, Microsoft).

### Scanning electron microscopy (SEM)

Colonies from the 2012 and 2013 embryo dish experiments were placed in a freezer at −20°C in 2.5 mL of 0.2 μm filtered, oxic seawater for 9.5 min prior to fixation to avoid contraction of the colonies [21]. Before the freezing point was reached, the embryo dish was taken out and 2.5 mL of modified Trump’s fixative (2.5 % glutaraldehyde, 2 % paraformaldehyde in 0.1 M sodium cacodylate buffer 1100 mOsm, pH 7.2, filtered with a 0.2 μm filter prior fixation) was added (modified from [33]). The samples were immediately rinsed with this solution and stored until further treatment.

After storage in fixative for a few months, colonies were rinsed in 0.1 M sodium cacodylate buffer (1100 mOsm, pH 7.2) three times for 3 min each, dehydrated in acetone and transferred to a mixture of acetone/hexamethyldisilazane (HDMS) (1:1) for 15 min, followed by two baths of pure HDMS for 15 min each. Subsequently, the samples were air dried for 3 h, placed on a stub and sputter coated with gold-palladium using an Agar Sputter Coater Agar 108 for 250 seconds.

For detailed SEM observations on a Philips XL 20 scanning electron microscope (acceleration voltage of 20kV) we used two sets of three colonies kept in oxic seawater for 48 h (the set from 2012 was used for statistical analyses, the other one from 2014 for additional SEM micrographs) and as control three colonies freshly collected from the environment in 2012. From each colony, images were taken from 15 microzooids (feeding cells) at a magnification of 2000x. The following symbiont parameters were analyzed for the oral and the aboral part of each microzooid separately: 1) number of symbiont cells in a 70 μm^2^ rectangular frame; all cells crossing the edge of the frame were counted only along one length and width of the frame; cells crossing the whole frame either longitudinally or horizontally were also counted. 2) The percentage of symbionts covering host surface (host coverage) was measured with Gimp 2.8 software after manual segmentation of the bacteria, whereby all partially and total cells in the 70 μm2 frame were included in the analysis. All other parameters were analyzed with AnalySIS^®^ program (Soft Imaging System GmbH, Münster, Germany), for each oral and aboral part of the microzooids up to 70 cells each, whereby cells were selected in a clockwise helical pattern: 3) length, 4) width, and 5) frequency of dividing cells (FDC). Dividing cells were defined as bacteria showing an invagination but not a clear intervening zone between cells [34]. 6) Cell volume was calculated from length and width data considering each cell as a cylinder plus two hemispheres [35]. 7) The cell elongation factor (EF), the ratio of length to width, was calculated for each cell [36]. The larger the EF the more rod-shaped the cells are, while cocci have an EF of approximately 1.

Statistical analyses were conducted with R [27] on data from three colonies kept at oxic conditions for 48 h and from three colonies collected *in situ* in 2012 (S2 Table). Because the Shapiro-Wilk tests performed for all parameters for each part of each microzooid showed deviations from normality, we used the Wilcoxon-Mann-Whitney test to evaluate differences within and between *in situ* and 48h oxic conditions.

### DNA extraction, PCR amplification and Sanger sequencing of incubation water

DNA was extracted from one half of each filter either fixed in ethanol or frozen (see above) using the FastDNA^®^ SPIN Kit for Soil (Bio 101, Carlsbad, CA) according to the manufacturer’s instructions. For DNA amplification, a nested-PCR approach was implemented. The first amplification was done using the universal bacterial primers 27 forward and 1492 reverse [37]. Following the first amplification, 1 μL of the PCR product was used as template for the second amplification using the primers 196 forward and 1439 reverse specific for the 16S rRNA of *Cand*. Thiobius zoothamnicola [24]. PCR amplifications were performed using a standard PCR cycling program (3 min at 95°C, 35 cycles of 30 sec at 95°C, 30 sec at annealing temperature, and 1 minute and 30 sec at 72°C, plus an additional 10-min cycle at 72°C) and an annealing temperature of 50°C for the universal bacterial primers and 58°C for the specific primers. As a positive control, DNA extracted from one colony collected in the harbor of Saint Bernardin (45°30’52.5”N, 13°34’24.3”E) was used. Sanger sequencing was performed on products extracted from seawater and from symbionts on one colony as a positive control.

## Results

### Host response to oxic conditions

*In situ* collections for investigating the host response to experimental oxic conditions came from two close by subtidal locations in the northern Adriatic Sea, the Strunjan mudflat and the Sv. Jernej estuarine. No significant differences were found in colony size (Wilcoxon-Mann-Whitney test: W = 941, p = 0.72, n = 85, S3A Fig) and in numbers of initial macrozooids on colonies between the two sites (W = 1931, p = 0.49, n = 120, S3B Fig). Although the number of swarmers released during oxic conditions was highly variable and ranged from 0 to 21 swarmers per colony, both populations did not differ significantly (W = 1936, p = 0.46, n = 120, S3C Fig). Also swarmer size was not significantly different between populations (W = 1306, p = 0.41, n = 33, S3D Fig). Therefore, we pooled the data from both locations for further analyses.

The mortality of the colonies showed a sigmoid pattern (bGLM: D^2^ = 0.93), strongly increasing 24 h after the start of the experiment and showing progressively smaller increases after 60 h (Fig. 1A). All times are expressed as the upper bounds of the observation intervals. A LT_50_ of 44 h (estimated standard error SE = 11 h) was calculated. The median colony survival time was 48 h (interquartile range IQR from 36 h to 60 h, n = 120). A maximal survival time of 96 h could be observed in two out of 120 colonies.

**Fig 1.**
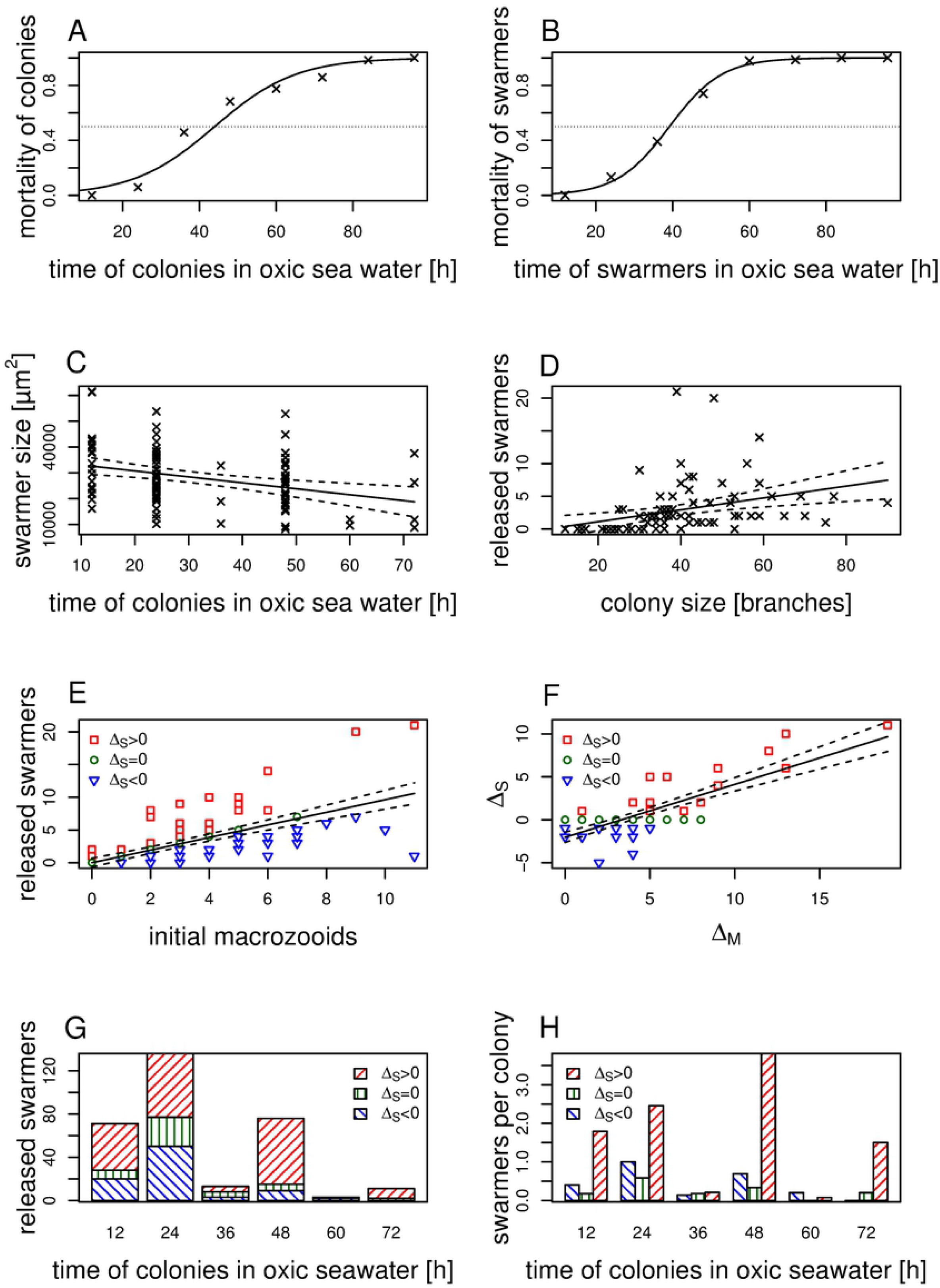
Host response to oxic conditions. (A) Binomial Generalized Linear Model of the mortality of the colonies given as the proportion of dead colonies in relation to the total number of colonies. LT_50_ is indicated as the point of intersection with the dashed line. (B) Binomial Generalized Linear Model of the mortality of swarmers given as the proportion of dead swarmers in relation to the total number of swarmers. The x-axis marks the upper bound of the swarmer survival time after release from the colony. (C) Ordinary least squares regression model with a confidence interval (CI) of 95% showing a negative correlation between swarmer size and time colonies spent under oxic conditions before releasing swarmers. (D) Positive correlation between colony size and the number of released swarmers. (E) The number of swarmers (released macrozooids) is positively correlated with the initial number of macrozooids. Δ_S_ is defined as the difference between the number of released swarmers and the initial number of macrozooids. Positive values of Δ_S_ indicate the net number of additionally released swarmers, whereas negative values display the net number of macrozooids remaining on the colony. Colonies are divided in three groups according to their production and release of swarmers (Δ_S_), positive Δ_S_ values are indicated in red, zero Δ_S_ values in green and negative Δ_S_ values in blue. (F) Δ_S_, the net production and release of swarmers, is positively correlated with Δ_M_ the production of macrozooids. (G) The swarmers were released in three cohorts with intervals of 24 h. (H) Colonies with Δ_S_ > 0 released more swarmers per colony and during longer time.

The mortality of swarmers displayed a similar pattern to the one of the colonies (bGLM: D^2^ = 0.99). The calculated LT_50_ was 39 h (SE = 9 h). The median swarmer survival time was 48 h (IQR from 36 h to 60 h, n = 143). The maximal life span of swarmers was 84 h (Fig 1B). Swarmer size strongly varied between individuals (median = 26601 μm^2^, IQR from 20737 to 35947 μm^2^, n = 99). Size measurements in 12 h intervals according to the time a swarmer spent in the oxic water, revealed no significant decrease (OLS: p = 0.08, F = 3.2, n = 99). Swarmer size, however, decreased in relation to the time colonies spent under oxic conditions (OLS: r^2^ = 0.11, p < 0.001, F = 12.2, n = 99, Fig 1C). Thus, colonies, which spent more time without sulfide, released significantly smaller swarmers.

The median size of the colonies was 43 branches (IQR from 38 to 53, n = 85) with a median number of initial macrozooids per colony of 2 (IQR from 0 to 4, n = 120). An OLS-regression model showed a linear relationship between colony size and initial number of macrozooids present on the colony prior start of the experiment (OLS: r^2^ = 0.20, p < 0.001, F = 20.7, n = 85, S4A Fig). Altogether, 310 swarmers were released (median number per colony 1, IQR from 0 to 4, n = 120). Colony size positively correlated with the number of released swarmers (OLS: r^2^ = 0.12, p = 0.001, F = 10.9, n = 85, Fig 1D). The number of swarmers released per colony, did not significantly correlate with colony longevity (OLS: p = 0.10, F = 2.8, n = 120).

To investigate whether the released swarmers came from macrozooids that were present already prior start of the experiment or from macrozooids that developed during the experiment we calculated the production of new macrozooids (Δ_M_) under oxic conditions. Nine negative values of Δ_M_ (and their corresponding Δ_S_) were discarded in subsequent statistical analyses, due to initial macrozooids which final destination remained unaccounted for. Δ_M_ strongly differed among colonies with a median of 3 (IQR from 1 to 5), ranging from 0 with no additionally produced macrozooids to maximal 19 additionally produced macrozooids. Colonies carrying a higher number of initial macrozooids produced a higher number of macrozooids (Δ_M_) under oxic conditions (OLS: r^2^ = 0.20, p < 0.001, F = 17.0, n = 71, S4B Fig).

A positive correlation between the number of initial macrozooids and the number of released swarmers was found (OLS: r^2^ = 0.47, p < 0.001, F = 106, n = 120, Fig 1E), but the number of initial macrozooids and unreleased macrozooids still on the colony at the end of the experiment did only weakly correlate (OLS: r^2^ = 0.06, p = 0.03, F = 5.0, n = 80, S4C Fig). The median number of unreleased macrozooids at the end of the experiment was 3 (IQR from 1 to 5, n = 80). To investigate whether colonies only released macrozooids that were present already prior start of the experiment or were able to release additionally produced macrozooids (Δ_M_), we calculated Δ_S_, the number of produced macrozooids that were also released as swarmers during oxic conditions. Δ_S_ ranged from −5 to 11 with a median of 0 (IQR from −1 to 0, n = 71). This indicates that in some colonies reproduction continued under oxic conditions (Δ_S_ > 0, n = 17), whereas other colonies produced and released no additionally swarmers and none (Δ_S_ = 0, n = 31) or only few initially present macrozooids were released into the water but died on the colony (Δ_S_ < 0, n = 23). Swarmer size showed no obvious relation with the production and release of swarmers Δ_S_ (OLS: p=0.07, F = 3.2, n = 99). There was a positive correlation between the produced macrozooids (Δ_M_) and the produced and released swarmers (Δ_S_). According to its slope, the production of ten additional macrozooids was needed in order to effectively release six additional swarmers (Δ_S_ = −2.0 + 0.6 * Δ_M_, r^2^ = 0.63, p < 0.01, F = 120, n = 71, nine colonies with Δ_M_ < 0 excluded, Fig. 1F). Swarmer release through time followed three consecutive cohorts in 24 h intervals with more than half of all swarmers leaving the colonies during the first day (Fig. 1G). Longer-lived colonies with positive Δ_S_ showed increased individual performance in the second swarmer cohort after 48 h in oxic water (Fig. 1H). Interestingly, the colonies with positive Δ_S_ values were of intermediate sizes (S4D Fig) and showed a higher longevity with a LT_50_ of 56 h (SE = 10 h).

### Symbiont response to oxic conditions

To investigate the change in symbiont coverage and colonization of other microbes in relation to host survival and time we repeated the oxic experiment with 10 to 20 colonies in different embryo dishes. We performed FISH using a symbiont-specific probe and a mix of archaea and bacterial probes on semi-thin sections of a few selected colonies removed after each time point. All hosts survived for up to 24 h. At this time point, the symbiont monolayer remained undisturbed in three out of four colonies, similar in appearance to three studied colonies immediately fixed after collection from the field. The fourth colony experienced some minor loss of symbionts.

There was little difference regarding the symbiont coverage on live and dead hosts after 48 h (n = 8; four live and four dead colonies) and after 72 h (n = 6; three live and three dead colonies), respectively. After 48 h, two out of the surviving four colonies showed little disturbance, i.e. little change in symbiont coverage (S5 A-D Figs), one colony showed major loss of symbionts, and one colony was aposymbiotic. Symbiont coverage on three out of four dead colonies was little disturbed, and the fourth dead colony was highly disturbed, i.e. major loss of symbionts. After 72 h two out of the three live colonies were aposymbiotic (S5 E-H Figs), the third colony was highly disturbed showing very few symbionts left. All three dead hosts were aposymbiotic.

The onset of epigrowth of other microbes, including bacteria and archaea, happened already within the first 24 h. We noted that microbial fouling mainly proceeded from the bottom part of the colony (S5 A-D Figs), also in part overgrown by microbes in nature [11,12]. Mostly, other microbes colonized host surfaces directly once the symbiont was lost (Figs 2 A-D). In some cases, however, the symbiont monolayer was overgrown directly (Figs 2 I-L). Unspecific epigrowth was found in all colonies regardless of viability of the host.

**Fig 2.**
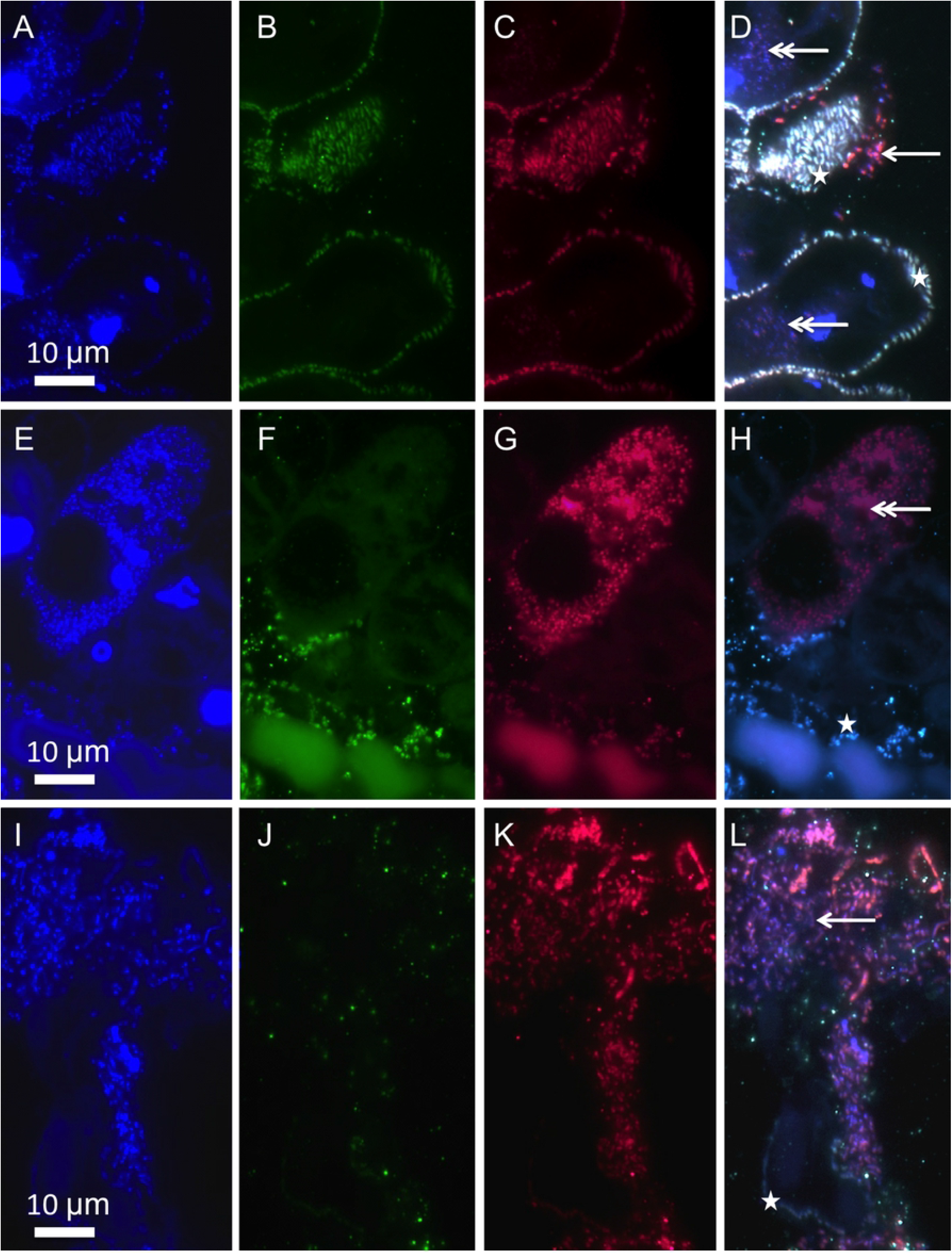
Symbiont response to oxic conditions applying FISH. Symbionts (asterisk), other epibiotic microbes (arrow), and intracellular microbes in D confined to small areas most likely food vacuoles, in H filling the entire host cell most likely infection (double arrow); (A, E, I) DAPI staining (blue), (B, F, J) symbiont-specific probe (green); (C, G, K) EUB_mix_ and Archaea probes (red); (D, H, L) composite of DAPI, symbiont-specific and EUB_mix_/Archaea probes. A-D and I-L from live colony after 48 h, E-H from dead colony after 48 h.

In addition, in some individual microzooids unidentified microbes were detected. Some were confined to small patches and were most likely contained in food vacuoles (Figs 2 A-D). Others, however, were filling the entire host cell, we interpret as potential microbial infection (Figs 2 E-H). These individual host cell infections appeared to increase in number with time of incubation and were dispersed randomly within the colony. Because we did not find branches, stalks, or clustered groups of infected microzooids, the infection apparently did not spread from one infected to adjacent microzooids.

To account for the differences in symbiont morphology on the microzooids using SEM [12, 21], we distinguished between symbiont populations located on the oral and aboral part of the microzooids freshly collected from the environment (Fig 3A) and compared them to those kept under oxic conditions for 48 h (Figs 3 B-G). Further, parts of the colony became covered with a mucus-like substance (Fig 3B) and/or other microbes (Figs 3 F, G) consistent with the results obtained by FISH.

**Fig 3.**
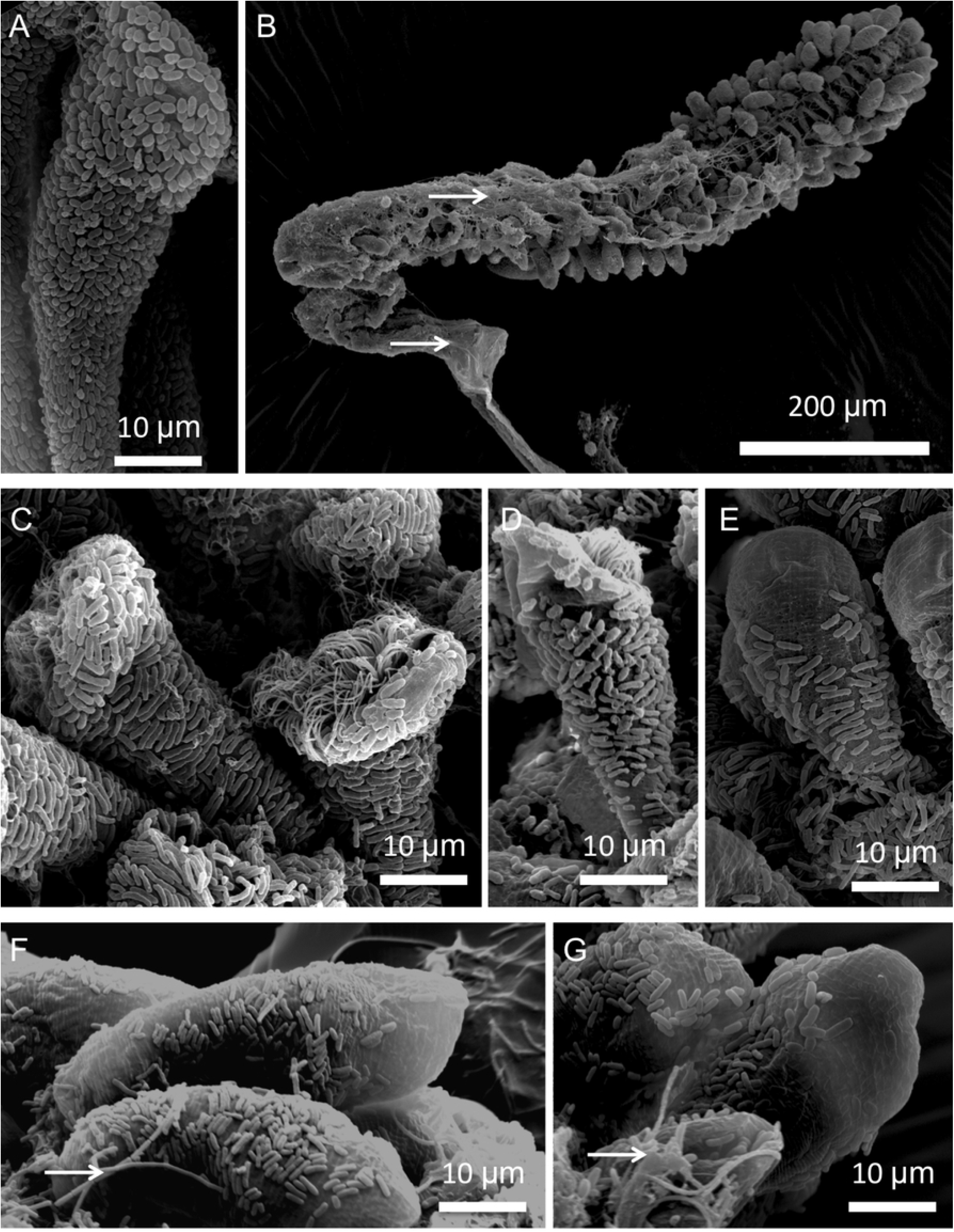
Symbiont response to oxic conditions applying SEM. Microzooid from colony prior experiment (A), and several colonies after 48 h in oxic seawater (B-G); (B) overview of colony covered in part with mucus; (C-G) microzooids with symbionts fully covering the host, and with gradually less and less coverage; arrows point to very long rods most likely not symbionts.

In freshly collected colonies (n = 3, Fig 3A), the microzooids were covered with a monolayer of symbionts, with similar symbiont coverage values in the oral and in the aboral part (Table 1). The host exhibited significantly higher numbers of symbionts per unit of surface on the aboral part than on the oral part of microzooids (Table 1). Orally located symbionts were significantly longer and wider than on the aboral part; hence orally located symbionts had a larger volume than aborally located ones (Table 1). The cell elongation factor, calculated as the length divided by the width, however, showed that aboral located symbionts were more rod-shaped than orally located ones (Table 1). The FDC of the symbiont population at the oral part was significantly higher than that at the aboral part (Table 1).

**Table 1.**
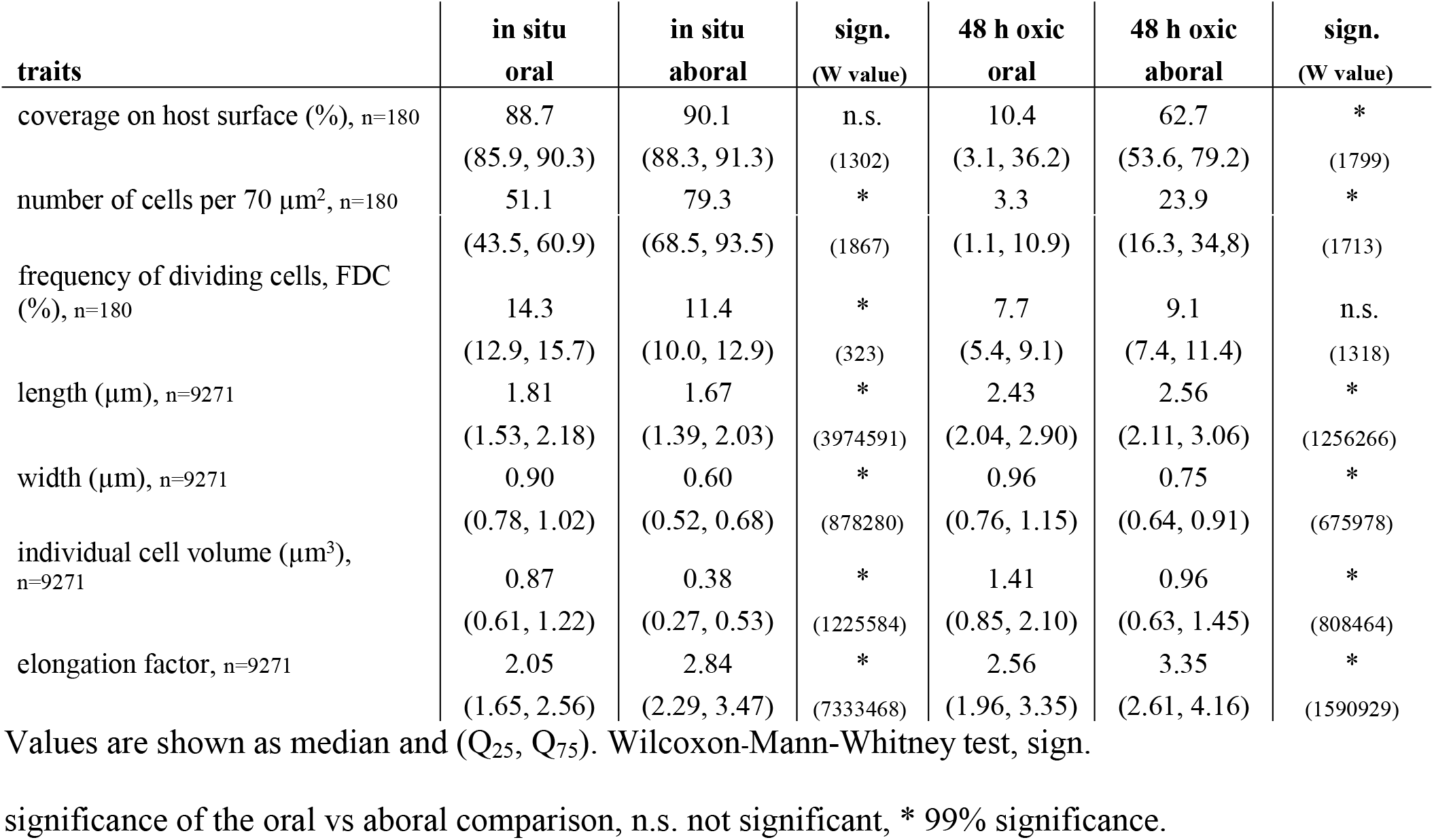
SEM analyses of symbiont traits.

After 48 h in oxic seawater the oral and the aboral symbiont populations were significantly different at the 1% level of significance in all parameters when compared to freshly collected colonies (Table 1). Orally, symbiont coverage was very low with a few symbionts remaining, compared to a higher coverage aborally (Table 1). Symbionts were still longer and wider orally than aborally, having a higher cell volume and a lower elongation factor (Table 1).

Overall symbiont coverage on both parts of the microzooids changed dramatically within 48 h in oxic seawater. A significantly lower symbiont coverage (Figs 4 A, B) and symbiont number on both oral and aboral parts of the microzooids (Figs 4 C, D) were observed compared to freshly collected colonies. Symbionts on both parts of the microzooids significantly increased in volume (Figs 4 E, F) and became significantly more rod-shaped (Figs 4 G, H). Also, FDC was significantly lower after 48 h (Table 1, Figs 4 I, J) compared to freshly collected colonies.

**Fig 4.**
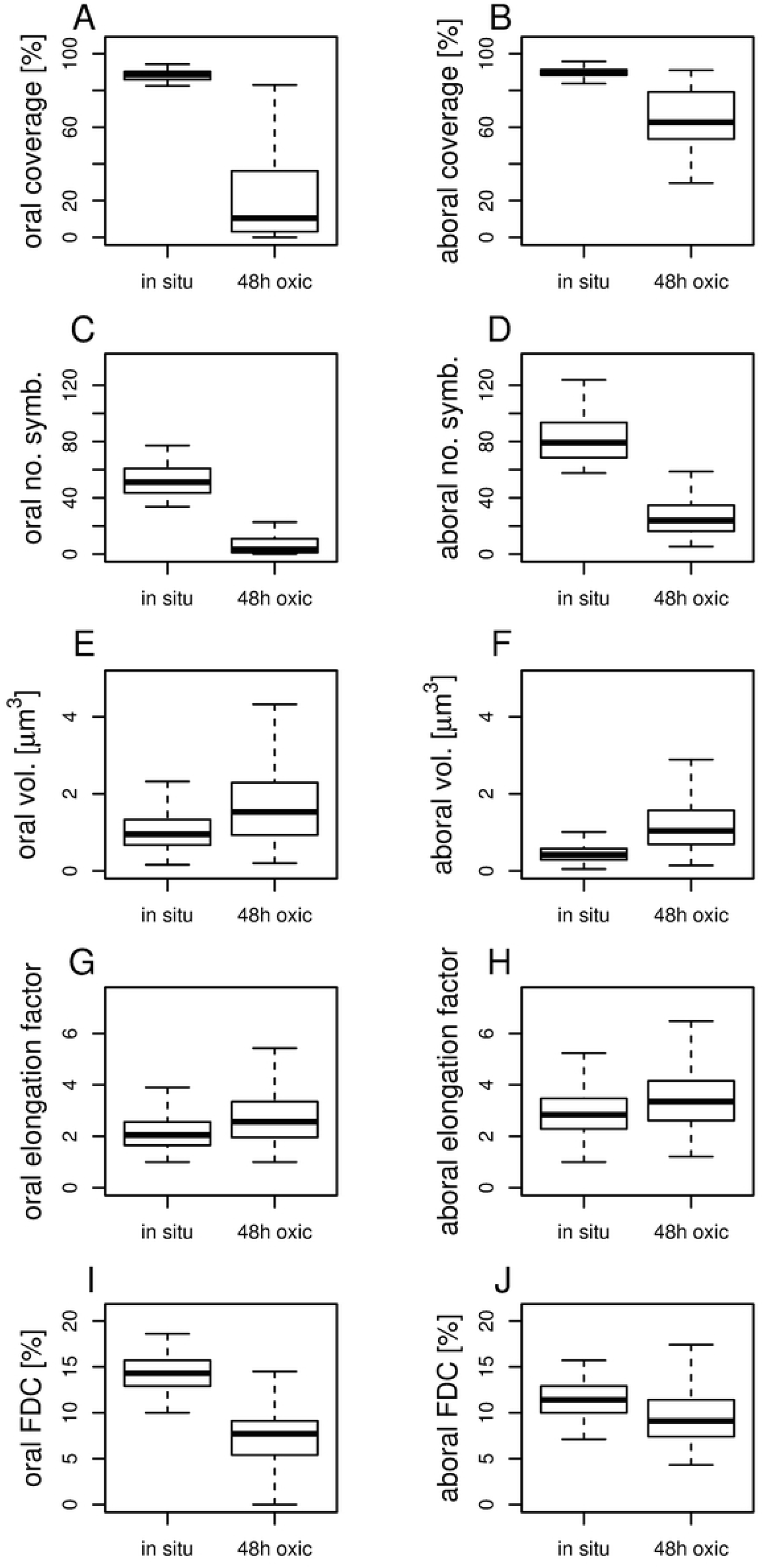
Symbiont response to oxic conditions compared to freshly collected colonies. Box-and-wisker plots comparing orally (A, C, E, G, I) and aborally (B, D, F, H, J) located symbionts on microzooids of following parameters: percentage of symbiont coverage on host outer surface (A, B), number of symbionts per 70 μm2 host surface area (C, D), volume of individual symbiont cells (E, F), cell elongation factor (ratio of symbiont length to width) (G, H), and frequency of dividing cells (FDC, ratio of dividing to total symbiont cells) (I, J). All data were compared with the Wilcoxon-Mann-Whitney test and proved significantly different (99% significance) between *in situ* conditions and 48 h oxic experiments.

The nested-PCR amplification using universal bacterial and subsequently symbiont-specific primers revealed the presence of the 16S rRNA symbiont phylotype in some of the experimental incubation waters. Positive results were obtained from seawater of one embryo dish already after 12 h of oxic incubation, and thereafter also after 24 h and 48 h (whereby the incubation water was exchanged after each time point). Water from the other tested embryo dish was only positive after 72 h of oxic treatment.

## Discussion

While maintenance of host-microbe mutualism over a host generation requires finely tuned exchange of benefits between partners, persistence over ecological time scales requires reproduction prior host death and transmission of symbionts from one to next host generation [14, 38, 39]. In unstable environments such as those inhabited by the giant ciliate mutualism, one of the biggest, naturally occurring threats is the cessation of sulfide flux. We have shown in a suite of experiments that the association breaks down quickly when exposed to such oxic conditions. Host and symbiont, however, reacted differently. Host colony reproduction through swarmers was maintained until host death in less than two days. Symbiont morphology on the host changed and division was reduced within two days compared to *in situ* values. In addition, some symbionts left the stressed and/or dead hosts back into the environment.

Our experiments show that host reproduction under oxic conditions continued until host death. Swarmers not only came from macrozooids that were present on the colony already prior start of the experiment but also from macrozooids that developed during the experiment. Because not all initially present macrozooids and not all newly developed macrozooids were released, overall the number of released swarmers roughly matches the number of initial macrozooids present. This clearly points to the importance of macrozooid production until host death contributing to host net reproduction even when environmental conditions become detrimental for the association.

Altogether, 310 swarmers were released from 120 colonies, with more than half of all swarmers leaving the colonies in the first day. Release did not cease until host death, with swarmer size decreasing significantly with time the colony spent under oxic conditions. This indicates a trade-off between quality and quantity of offspring following the strategy of *r*-selection, common for species from unstable environments [40].

We suggest that lack of sulfide resulted in reduced nourishment of the host. Prior studies showed that under prolonged oxic conditions carbon fixation in the symbiont ceases and subsequently release and uptake of organic carbon also stops [28]. Symbionts on colonies kept under oxic conditions for 24 h prior ^14^C or ^13^C labeled bicarbonate incubations showed no carbon fixation and incorporation and no uptake into host tissue occurred [28]. Host nourishment under such oxic conditions then is reduced to direct feeding on free-living microbes and symbionts [28]. We do not know yet, whether the changes in host nutrition alone or other yet unknown benefits not provided by the symbiont under oxic conditions led to a stressed host that died in less than two days.

A remarkable phenotypic change happened in the symbionts under oxic conditions in as little as two days. Differences in morphology between symbionts on oral and aboral microzooid parts, known to be present in freshly collected colonies from the environment [21], were maintained, but symbionts on both parts became more rod-shaped and grew larger compared to freshly collected colonies.

Surprisingly, FDC values indicate that proliferation did not completely cease as expected, but was highly reduced. Given the fact that internal sulfur storage in the symbionts can only support carbon fixation for a very short time [23,28], the symbiont may switch to heterotrophic metabolism and therefore sustain proliferation. Although genes supporting this function are present in the metagenome assembled genome (Espada-Hinojosa unpubl.), whether they are expressed under such conditions remains to be studied. FDC values were not significantly different (at a 1% level of significance) in orally and aborally located symbionts exposed to oxic seawater (Table 1). This points to more similar abiotic conditions for the symbionts regardless of location on the microzooids, consistent to previous cultivation experiments under a steady flow of oxic seawater but supplemented with sulfide where oral and aboral symbionts also exhibited similar FDC values [21]. We suggest that under the oxic conditions lacking reduced sulfur species we applied here the symbionts responded with similar proliferation behavior on all parts of the microzooids possibly due to heterotrophic uptake, while symbionts exposed to a steady flow of sulfide and oxygen accordingly proliferated due to chemoautotrophy [21]. In contrast, orally located symbionts from colonies freshly collected from wood exhibited higher FDC values than aborally located ones, confirming earlier results from colonies collected from degrading seagrass leaves and from vertical, overhanging rocks above seagrass debris [21].

Host–symbiont maintenance was clearly disturbed under oxic conditions. The symbiont coverage on the host was significantly reduced compared to freshly collected colonies exhibiting a monolayer on the host. This may have resulted from reduced symbiont proliferation under oxic conditions combined with loss of symbionts due to host ingestion. Further, finding the symbiont phylotype in the incubation water of some of the experiments strongly points to loss of symbionts from the host surface either due to host release or symbiont escape into the environment. Whether these symbionts, however, are viable to colonize other hosts following a wait-and-sit strategy known in some pathogens [41] or even to proliferate as a free-living population, is not known.

Disturbance of host – symbiont maintenance was also evident by microbial fouling on symbiont-free host surfaces or even on top of the symbiont. In freshly collected colonies epigrowth proceeds from the bottom part of colonies [11, 12] similar to our observations of stressed hosts. Because these are the oldest parts of the colony this may indicate that host age plays a role in defense against microbial fouling under natural sulfidic as well as experimental oxic conditions. Alternatively, or in addition, symbionts may contribute to the antimicrobial defense.

Not only the numbers of swarmers produced per colony, also survival of the swarmer is crucial because it sets the limits of dispersal to find a patchy, sulfide-leaking habitat for settlement. With a swarmer LT_50_ of 39 h and a swim speed of 5 mm s^-1^ [42] dispersal over about 700 m can be accomplished given the swarmer swims in a straight line. Because the vertically transmitted symbiont on the swarmer [11,12], however, is lost under oxic conditions within 24 h [25], the maximal estimated swarmer dispersal distance can only be reached together with the symbiont if the swarmer migrates in the sulfidic boundary layer close to sulfide emitting surfaces. In oxic seawater, symbiotic swarmers only reach about 400 m distance and from there on the host travels aposymbiotically until it settles or dies.

In conclusion, the beneficial interactions between *Zoothamnium niveum* and its sole symbiont *Cand*. Thiobius zoothamnicola become rapidly disturbed under stressful oxic conditions lacking sulfide. Colonies die quickly in less than two days but continue to reproduce until death releasing their asexually produced offspring to search for a suitable habitat. Symbionts are also affected rapidly, change their morphology and decelerate division. Now that the principal mode of stress response is known we can start to decipher the underlying mechanisms of changes in physiology and interactions on molecular level.

## Supporting Information

**S1 Fig. Time schedule of the experiment.** Colonies (n = 120) were monitored every 12 h (horizontal time line). Released swarmers from each of this time points were divided in 4 cohorts (A, B, C, D; vertical time line).

(TIFF)

**S2 Fig. Scheme of colony.** Colony composed of stalk and alternate branches and three different cell types – terminal zooids, microzooids, and macrozooids; size of colony counted in number of branches; colony with initial number of macrozooids present prior experiment and remaining number of macrozooids present at the end of experiment; during experimental time the release of swarmers was also counted. (TIFF)

**S3 Fig. Comparison of colonies from Sv. Jernej and Strunjan**. All Wilcoxon-Mann-Whitney tests fail to reject the null hypothesis of equal medians: (A) colony size (p = 0.72), (B) number of initial macrozooids per colony (p = 0.49), (C) number of swarmers (released macrozooids) per colony (p = 0.46) and (D) size of swarmers (p = 0.41).

(TIFF)

**S4 Fig. Additional aspects of host response to oxic non-sulfidic conditions**. (A) Colony size shows a positive correlation with the initial number of macrozooids in the colonies at the beginning of the experiment. (B) The initial number of macrozooids also positively correlates with the number of newly produced macrozooids (Δ_M_). (C) The initial number of macrozooids on the colonies and the final number of macrozooids that were not released as swarmers at the end of the experiment are positively correlated, although only with a weak significance at the 5% level. (D) Colonies that were particularly proficient in producing and releasing swarmers (Δ_S_ > 0) showed intermediate sizes.

(TIFF)

**S5 Fig. FISH micrographs colonies**. Colony alive after 48 h (A) DAPI staining (blue), (B) symbiont-specific probe (green), (C) EUB_mix_ and Archea probes (red) (D) composite of A, B, C; note the increase in microbial fouling from top to bottom. Colony alive after 72 h with very few symbionts left (E) DAPI staining (blue), (F) symbiont-specific probe (green), (G) EUB_mix_ and Archea probes (red), (H) composite of F, G and H.

(TIFF)

**S1 Table. Collections and abiotic parameters measured prior collection at wood surface.** Samples listed according to type of experiments, applied techniques, and time series of experiment, site, date of collection and number of wood, and abiotic parameters depth, temperature, salinity, and pH.

**S2 Table. Experiments and abiotic parameters measured at the start (and at the end) of experiment.** Abiotic parameters: temperature, salinity, pH, and percentage of oxygen saturation (mean ± standard deviation).

(DOCX)

## Acknowledgements

We would like to acknowledge the Marine Biology Station Piran (Slovenia) for their hospitality. EM work was performed at the Core Facility Cell Imaging and Ultrastructure Research, University of Vienna.

## Author Contribution

Funding acquisition, supervision, project administration: MB.

Formal analysis, investigation, methodology: MB. JD. SEH. JK. EK. IP. LS. JMV.

HCZ.

Visualization: JD. JK. EK. IP. LS. HCZ.

Writing of original draft: MB. SEH.

Review and editing of draft: MB. EK. SEH. LS. JMV.

